# The impact of chromatin dynamics on Cas9-mediated genome editing in human cells

**DOI:** 10.1101/071464

**Authors:** René M. Daer, Josh P. Cutts, David A. Brafman, Karmella A. Haynes

**Affiliations:** Arizona State University, School of Biological and Health Systems Engineering, 501 E. Tyler Mall, ECG 344A, Tempe, AZ, USA; Arizona State University, Biological Design Graduate Program, Tempe, AZ, USA 85287

**Keywords:** Cas9-mediated editing, chromatin, INDEL

## Abstract

In order to efficiently edit eukaryotic genomes, it is critical to test the impact of chromatin dynamics on CRISPR/Cas9 function and develop strategies to adapt the system to eukaryotic contexts. So far, research has extensively characterized the relationship between the CRISPR endonuclease Cas9 and the composition of the RNADNA duplex that mediates the system’s precision. Evidence suggests that chromatin modifications and DNA packaging can block eukaryotic genome editing by custom-built DNA endonucleases like Cas9; however, the underlying mechanism of Cas9 inhibition is unclear. Here, we demonstrate that closed, gene-silencing-associated chromatin is a mechanism for the interference of Cas9-mediated DNA editing. Our assays use a transgenic cell line with a drug-inducible switch to control chromatin states (open and closed) at a single genomic locus. We show that closed chromatin inhibits editing at specific target sites, and that artificial reversal of the silenced state restores editing efficiency. These results provide new insights to improve Cas9-mediated editing in human and other mammalian cells.

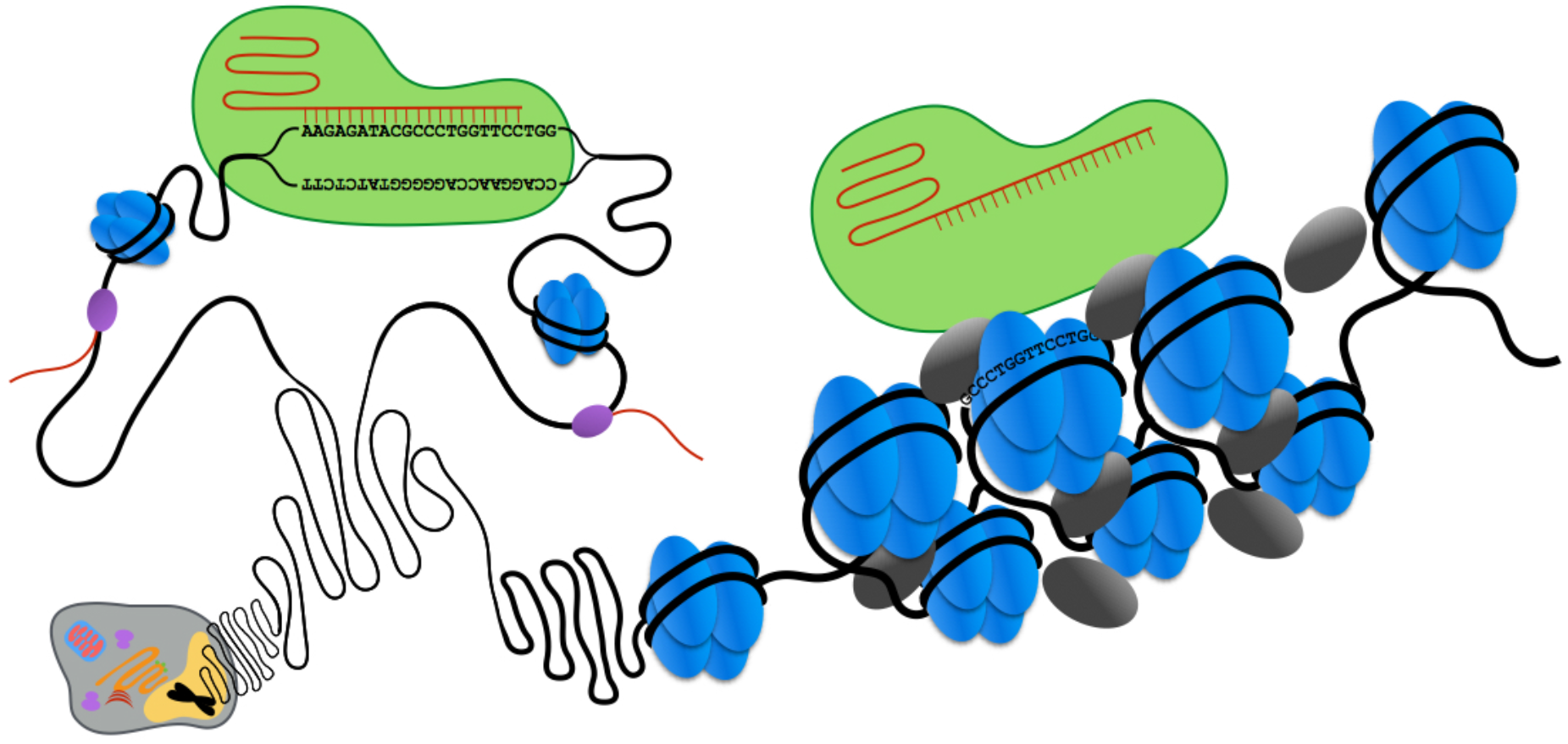

---

Extensive characterization and engineering is driving the CRISPR/Cas9 system to the forefront of biomedical research, human gene therapy, and tissue regeneration^1–4^. Realizing the full potential of CRISPR/Cas9 editing in eukaryotic cells will require efficient access to target sites within chromatin, the ubiquitous DNA-protein complexes that organize eukaryotic genomes, regulate gene expression, and render DNA less accessible to nucleases^5–7^. While some evidence suggests that chromatin modifications and DNA packaging can block eukaryotic genome editing by custom-built DNA endonucleases^8–11^, the underlying mechanism of Cas9 interference is unknown. Here, we present direct evidence that closed, gene-silencing-associated chromatin inhibits Cas9-mediated DNA editing. These results establish closed chromatin as a target for improving CRISPR/Cas9 efficiency in human and other eukaryotic cells.

There is conflicting evidence on whether chromatin interferes with Cas9 binding and nuclease activity in human cells. Chromatin immunoprecipitation (ChIP) mapping of dCas9, which lacks nuclease activity, showed preferential binding at off-target sites across the human genome with characteristics of open chromatin: high levels of DNase-hypersensitivity and protein-coding gene sequences^8,12,13^. Wu et al. (2014) reported reduced DNA cleavage at off-target sites near closed chromatin versus on-target sites near open chromatin^8^. While Cas9-mediated editing is not completely blocked at the silenced *SERPINB5* locus, which contains methylated DNA, the reported editing frequency was only ≤ 8%^9^. Taken together, these data suggest Cas9 may be inhibited by closed chromatin. However, Perez-Pinera et al. (2013) found that dCas9-based activators were functional at sites within closed chromatin, suggesting binding was not prevented^14^. So far, no study has compared Cas9 activity at a single site for both the open and closed chromatin state.

## Results and Discussion

Here, we use a model silencing system developed in Hansen et al. (2008)^15^ to control the chromatin state at a single site, a stably integrated *firefly luciferase* transgene. We directly measured the impact of closed chromatin on Cas9-mediated DNA editing by targeting Cas9 to sites within the *luciferase* gene in unsilenced, partially silenced, and fully silenced chromatin states. The Gal4EED HEK293 cell line contains a doxycycline (dox)-inducible transgene that expresses Gal4EED, which binds upstream of *luciferase* and recruits endogenous Polycomb Repressive Complexes (PRCs), key regulators of facultatively closed chromatin^16^ (Figure 1a, see Methods for detail). PRCs target hundreds of genes and play a critical role in gene silencing, embryonic development, stem cell maintenance and differentiation, and tumor suppression^17–21^, and thus, understanding whether differences in PRC accumulation influence Cas9 activity is necessary for advancing biomedical applications.

**Figure 1.**
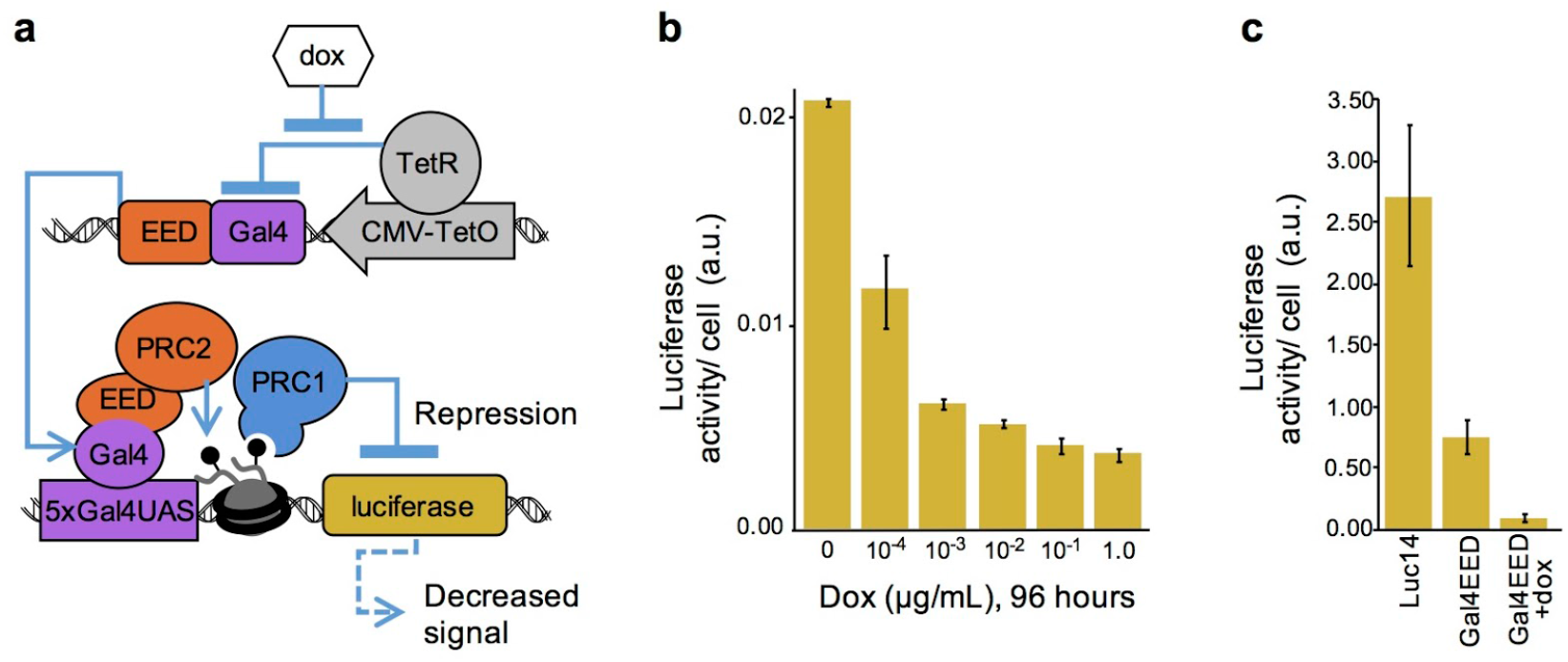
A closed chromatin state at luciferase is regulated by doxycycline (dox) in GAL4EED cells. (a) The GAL4EED circuit diagram illustrates how dox regulates the expression of the Gal4EED fusion protein, which mediates accumulation of Polycomb Repressive Complex 1 (PRC1) and closed chromatin at the *luciferase* reporter. (b) A repressed expression state at *luciferase* is stimulated by dox in GAL4EED cells. (c) Comparison of luciferase expression in cell that lack a *Gal4EED* gene (Luc14), and GAL4EED cells before and after treatment with dox. a.u.: arbitrary units from luminescence readings. Error bars indicate s.d. for n=3 technical replicates.

Gal4EED cells grown with 1 µg/mL dox for 96 hours show maximal levels of repression (Figure 1b). We observed that the Gal4EED cell line shows less *luciferase* expression than the parental Luc14 cell line, which lacks the *Gal4EED* transgene, perhaps due to leaky Gal4EED expression and a partially silenced chromatin state (Figure 1c). The chromatin states in GAL4EED are stable over time and in the presence of transfected plasmids. Once initiated by Gal4EED, the silenced state remains stable up to 96 hours after dox is removed and Gal4EED expression is no longer activated^15^. In our previous work, we showed that control plasmids that express fluorescent proteins do not alter the expression levels of active or Gal4EED-silenced *luciferase*^22^.

We constructed a series of plasmids expressing Cas9 and a short guide RNA (sgRNA) designed to target one of twenty-three sites within *Gal4UAS-TK-luciferase*. Editing efficiencies were determined using a SURVEYOR digestion assay and a Bioanalyzer (Figure 2a). The target sites are distributed across 900 basepairs (bp) of the transgene located downstream of the Gal4EED binding site (5xGal4UAS in Figure 2b). The sgRNAs showed a wide variability of INDEL frequency from zero (below detection limits) to almost 40% in Luc14 cells (Figure 2c).

**Figure 2.**
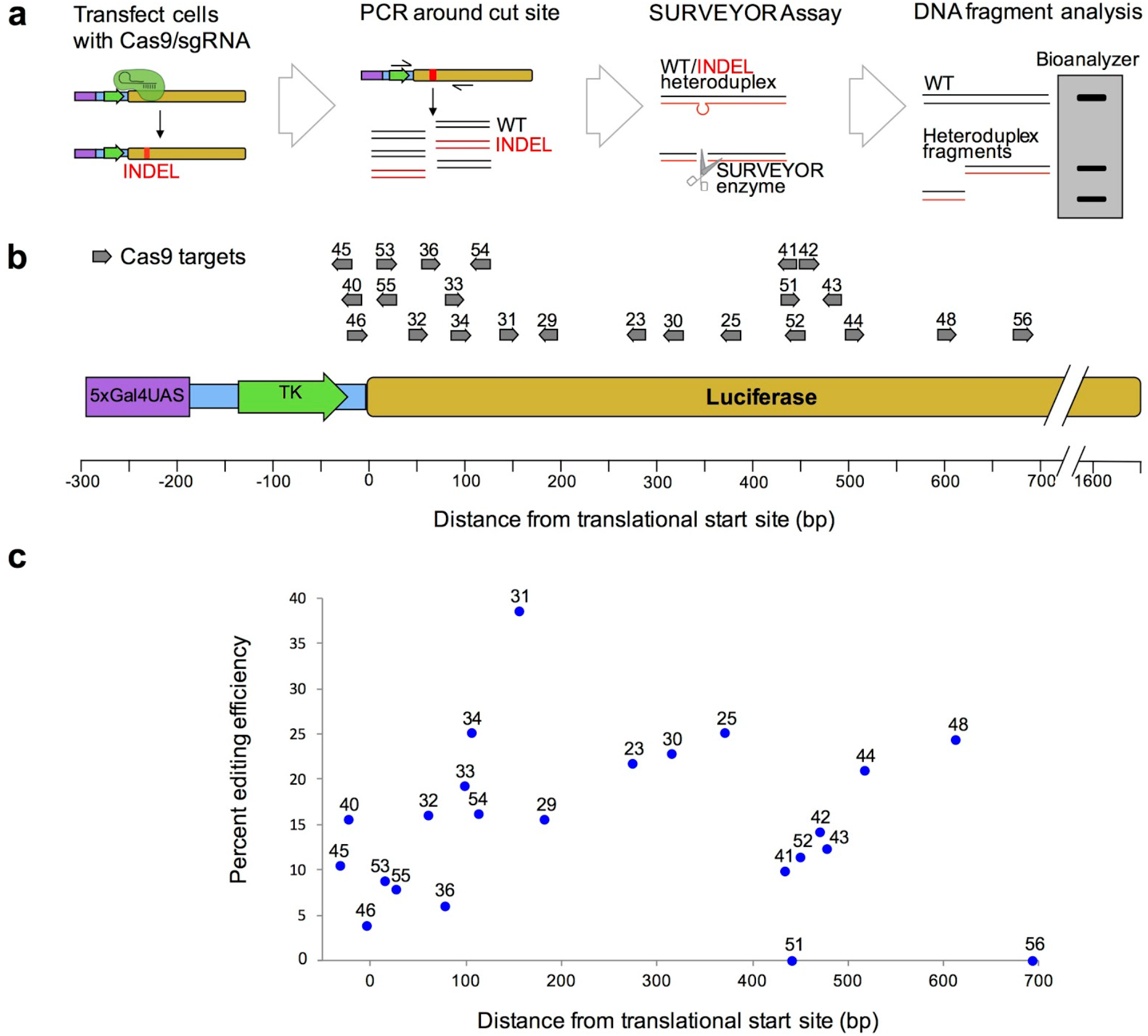
Editing efficiency at different Cas9/sgRNA target sites detected by SURVEYOR assays. (a) Diagram of the SURVEYOR assay. INDELS: Insertions and deletions; WT: wildtype (b) Map of all sgRNAs tested in this study. Arrows show Cas9/sgRNA binding sites. (c) SURVEYOR results from a screen to identify sgRNAs with detectable editing rates at the UAS-TK-luciferase reporter in Luc14 cells.

To determine the effect of facultative closed chromatin on Cas9-mediated editing, we compared the editing efficiencies of Cas9 in cells where the *luciferase* target was set to different expression states: unsilenced (Luc14), partially silenced (Gal4EED), and fully silenced (GAL4EED +dox) (Figure 1c). For this analysis, we used the five sgRNAs that showed the highest editing efficiencies in preliminary tests (Figure 2c) in Luc14 cells: sg034, sg031, sg025, sg044, and sg048. We also tested four additional sgRNAs, located farther upstream (sg046, sg055, sg032, sg054) in order to investigate Cas9 interference closer to the initiation site of chromatin compaction. In order to control for varying transfection and expression levels across cell lines and conditions, we added an enhanced green fluorescent protein (EGFP) reporter gene to the Cas9 plasmid (Figure 3a). In this plasmid, EGFP expression is driven by the same promoter as Cas9, but the T2A signal allows EGFP to be translated as a separate peptide to avoid interference with Cas9 function. We used the percentage of GFP-positive cells (Figure 3b) to normalize editing efficiency values across different experiments.

**Figure 3.**
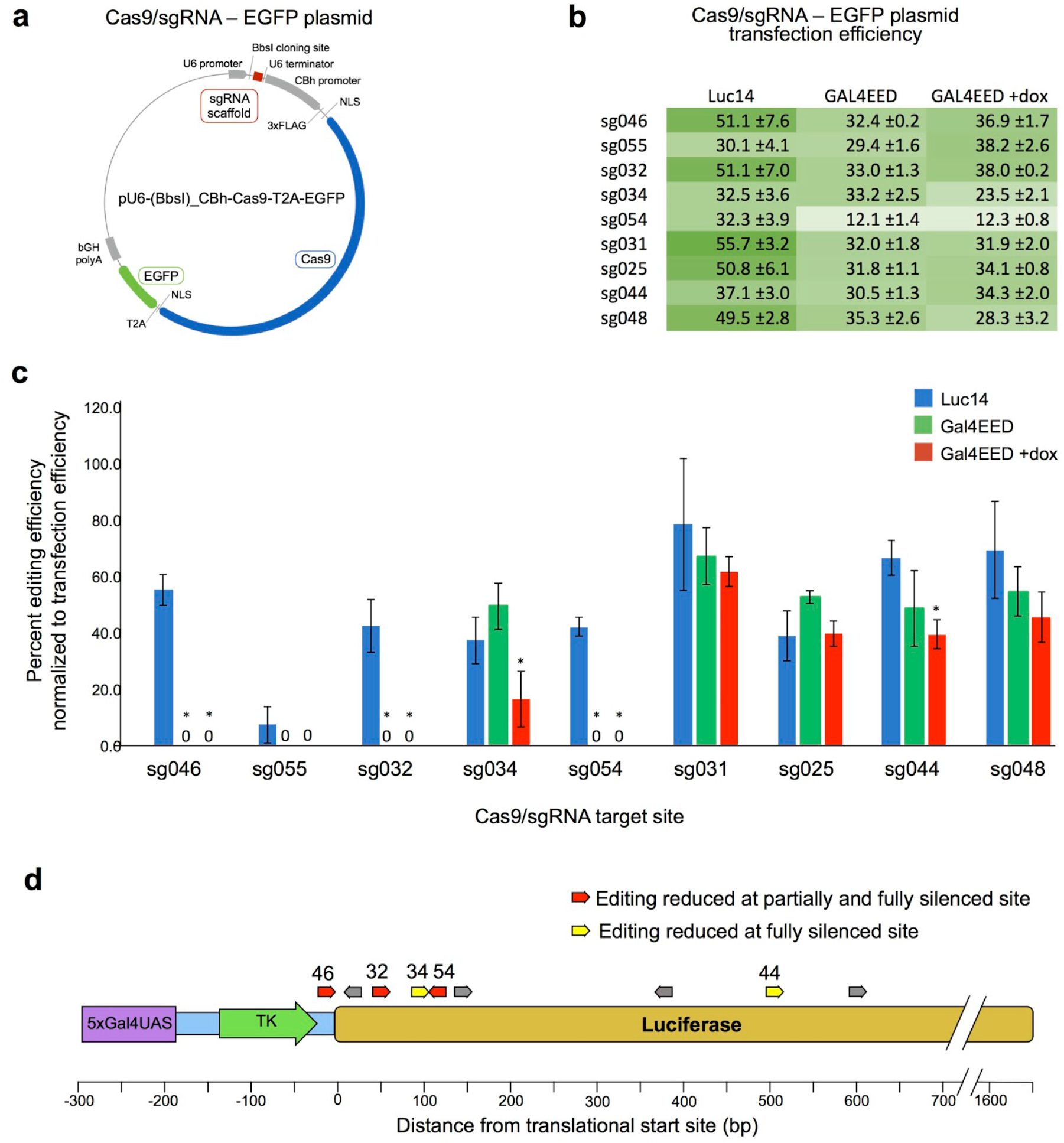
Cas9 editing efficiencies at target sites in unsilenced, partially silenced, and fully silenced chromatin states. (a) A map of the Cas9/sgRNA-expressing plasmid. Cas9 and EGFP expression are both driven by the CBh promoter. The T2A signal allows EGFP to be translated as a separate peptide to avoid interference of Cas9 function. (b) Table of average frequencies of EGFP-expressing cells (transfection efficiencies) as determined by flow cytometry of triplicate samples for transfected Luc14, GAL4EED, and GAL4EED cells treated with doxycycline. (c) Mean editing frequencies normalized to transfection efficiency in Luc14, GAL4EED, or GAL4EED +dox cells for Cas9 targeted to sites sg046, sg055, sg032, sg034, sg054, sg031, sg025, sg044, and sg048. * Indicates significantly reduced editing efficiencies at fully silenced chromatin compared to unsilenced chromatin (p < 0.025 for 3 biological replicates). Editing frequencies for target sites sg046, sg055, sg032, and sg054 for both GAL4EED or GAL4EED +dox cell types were below detection limits. Error bars indicate s.d. for n=3 biological replicates. (d) Summary of the data in (c). Cas9 target sites sg046, sg032, and sg054 show a reduction in editing efficiency in both the partially and fully silence states compared to the unsilenced states (red arrows). Cas9 target sites sg034 and sg044 show a reduction in editing efficiency in the fully silence states compared to the unsilenced states (yellow arrows).

Cas9-mediated editing was reduced at target sites in fully silenced chromatin compared to unsilenced chromatin for six of the nine sgRNAs we tested (Figure 3c). The greatest significant reductions occurred within 150 bp of the transcription start site downstream of the chromatin initiation site at 5xGal4 at sites sg046 (p = 0.002), sg055 (p = 0.092), sg032 (p = 0.008), and sg054 (p = 0.001). In the unsilenced state (Luc14), average editing efficiencies at these sites were 7.5 to 55.1% while mutation frequencies were below detection limits in both the partially (GAL4EED) and fully (GAL4EED +dox) silenced states. These results suggests that editing at TSS-proximal sites is sensitive to closed chromatin. Cas9-mediated editing in fully silenced chromatin was reduced, but not completely inhibited, farther downstream at sg034 (p = 0.024) and sg044 (p = 0.022) compared to unsilenced chromatin. Interestingly, editing efficiency at sg025, located between sg031 and sg044, and sg048, located downstream of g044, is not decreased in the presence of closed chromatin. This suggests that interference may occur in a Cas9/sgRNA-dependent manner or that the spreading of closed chromatin from the UAS is discontinuous. Overall, our results reveal differences in Cas9 accessibility at a greater resolution than what has been reported to date for a single genomic locus. Furthermore, comparison of open, moderately-closed, and fully-closed states at several on-target sites along a single locus allowed us to detect different levels of interference that are the direct outcome of the formation of facultative chromatin.

Next, we tested the hypothesis that in addition to reducing editing efficiency, silenced chromatin also affects the types of mutations that are generated at the target site. We analyzed mutations at target site sg034 because at this location the closed state still showed detectable Cas9-mediated editing. Sequencing of cloned mutants from Cas9-edited DNA confirmed lower editing efficiency at a target site in fully silenced chromatin compared to unsilenced chromatin. We then compared the distribution of mutated sequences from unsilenced- and fully silenced-chromatin samples. We detected various mutations that were generated in the absence of a repair template by non-homologous end joining (NHEJ) at target site sg034. These mutations primarily consisted of deletions ranging from 1 to 24 bp (Figure 4a). A small number of insertions (1 to 205 bp) and single base pair substitutions were also detected. DNA cloned from Cas9/sg034-treated *luciferase* in the unsilenced chromatin state showed a higher frequency and broader range of affected nucleotides compared to the fully silenced chromatin state (Figure 4b). The most frequent mutation was a single base pair deletion at the Cas9 cut site in both the unsilenced and fully silenced chromatin states (12.3% and 2.3%, respectively). This result led us to reject our hypothesis. The mutant library sequence data indicate that Cas9-mediated editing is reduced by ∼30% in fully silenced chromatin. SURVEYOR assays showed a similar reduction, ∼40%. Therefore, the sequencing data and the SURVEYOR data provide corroborating evidence that repressive chromatin interferes with Cas9-mediated editing at site sg034.

**Figure 4.**
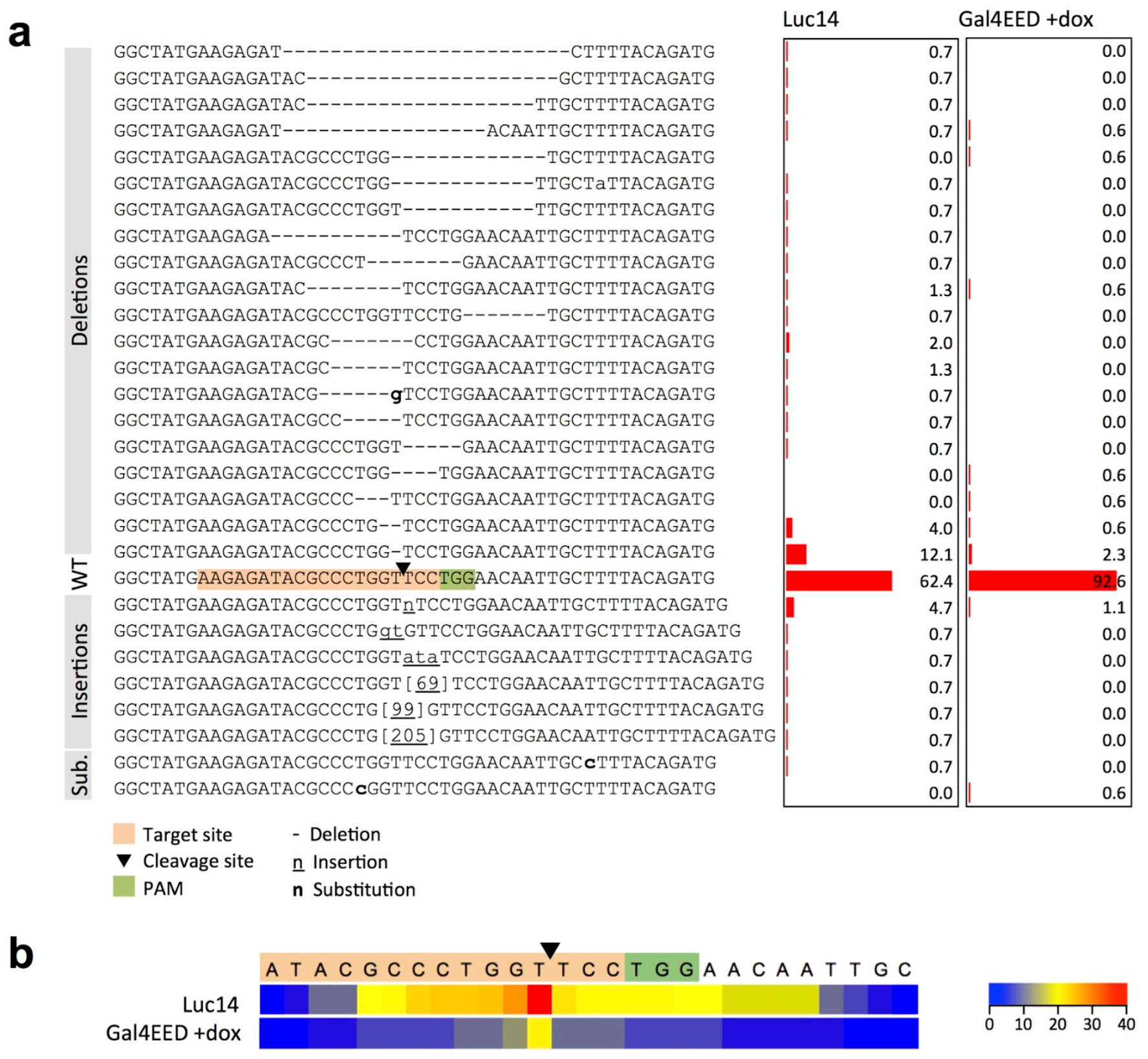
Frequencies of mutations in genomic DNA from Cas9/sg034-treated Luc14 or GAL4EED +dox cells. (a) Each row shows the mutated sequence aligned to the wild type sequence (WT). Bar graphs show the percentage of library clones that correspond to the sequence in the same row on the left. (b) The heat map shows the number of times each nucleotide position was affected by a deletion that arose from non-homologous end joining (NHEJ) repair.

In order to confirm the formation of repressive chromatin following dox treatment of GAL4EED cells, we performed chromatin immunoprecipitation (ChIP)-qPCR using an anti-H3K27me3 antibody (Figure 5a and Supplemental Figure S1). H3K27me3 is required for the maintenance of Polycomb-mediated chromatin compaction. We detected a 27- to 112-fold increase in H3K27me3 at *luciferase* in GAL4EED cells that were treated with dox for 96 hours. These data, along with our Luciferase expression assays in Figure 1b, demonstrate that we had induced facultative silenced chromatin at *luciferase.* Previous work by Hansen *et al.* showed through ChIP-qPCR that upon addition of dox, Polycomb Group 2 (PRC2) protein EZH2, Polycomb Group 1 (PRC1) protein CBX8, and the H3K27me3 mark accumulate at the *luciferase* transgene in GAL4EED cells^15^. We can conclude from their data that when Gal4EED protein binds upstream of *luciferase*, PRC2 is recruited and trimethylates H3K27. H3K27me3 binds PRC1, which is associated with nucleosome compaction and gene silencing.

**Figure 5.**
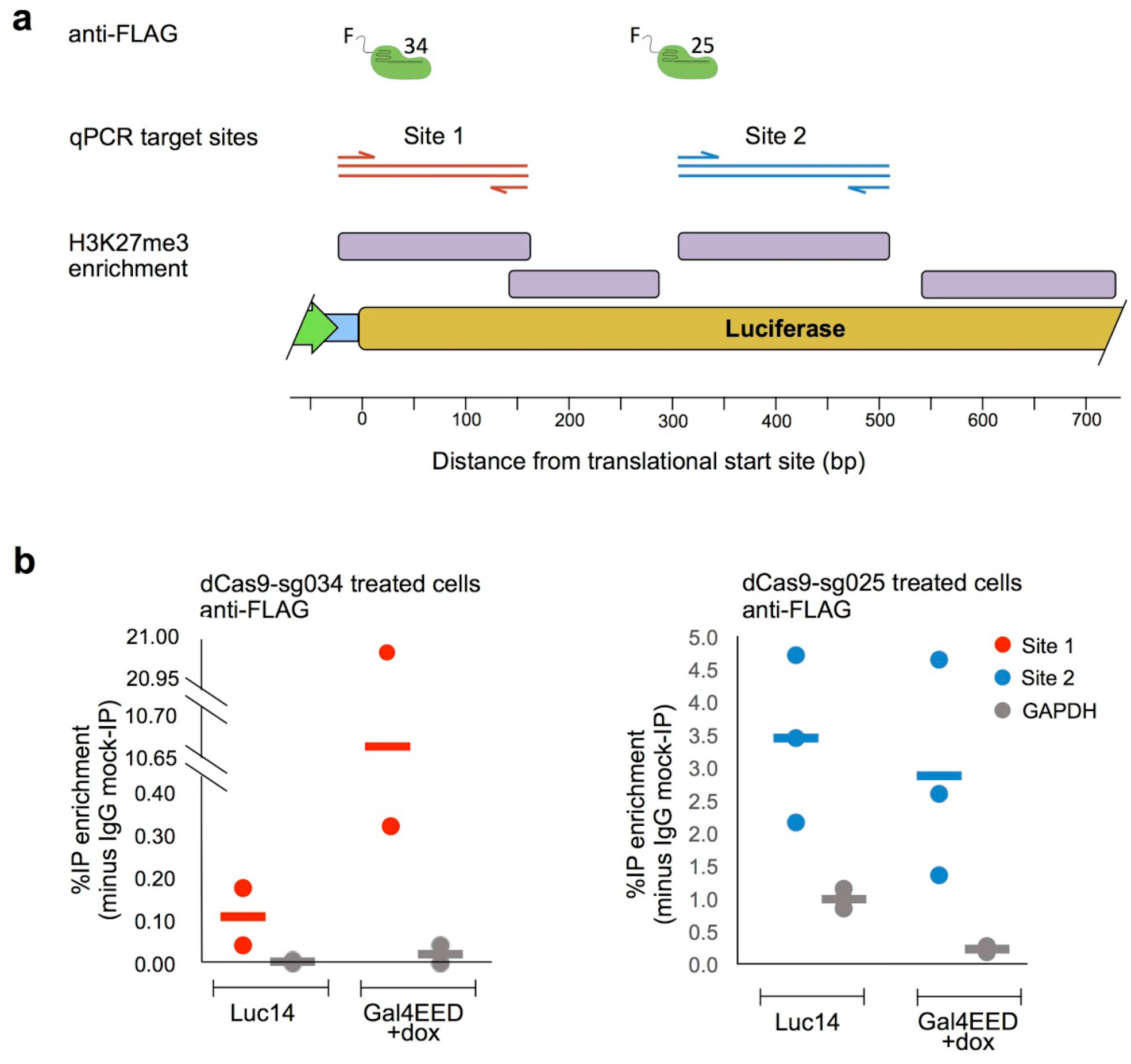
Chromatin mapping data show enrichment of H3K27me3 and dCas9 at gRNA target sites. (a) An anti-FLAG antibody or anti-H3K27me3 antibody was used to immunoprecipitate (IP) sheared, crosslinked chromatin from untreated Luc14 and GAL4EED cells. Quantitative PCR (qPCR) was used to measure IP’ed DNA. Primers (described in Methods) and amplicon sizes are shown in the illustrated maps. A primer pair located at the constitutively active GAPDH promoter was used as a negative control to determine off-target binding. H3K27me3 was enriched 27- to 112-fold in GAL4EED +dox cells compared to Luc14 cells (Supplemental Figure S1) at four sites spanning the Cas9/sgRNA target sites (purple bars). (b) Enrichments of IP DNA for two (dCas9/sg034-treated cells) and three (dCas9/sg025-treated cells) independent cross-linked chromatin replicates are shown as percentages of input minus background (IgG-IP DNA). Dots indicate individual measurements, bars indicate median (dCas9/sg034) or average (dCas9/sg025).

We performed chromatin mapping experiments to investigate whether interference of Cas9 activity is accompanied by a decrease in Cas9 binding. We used deactivated Cas9 (dCas9), which lacks DNA-cutting activity^23^, to analyze the binding activity of the Cas9/sgRNA complex at unsilenced and silenced chromatin. Formaldehyde-crosslinked chromatin was extracted from triplicate samples of dCas9/sgRNA-transfected cells and sheared to ∼700 bp. ChIP was carried out using antibodies against the FLAG tag at the C-terminus of dCas9. QPCR was used to determine the IP enrichment of target DNA sequences. We analyzed a site that showed Cas9 interference the the fully silenced chromatin state (sg034) and a site that showed no interference (sg025) (Figure 3c). We detected dCas9 enrichment at *luciferase* in both the open chromatin and closed chromatin states (Figure 5b). Taken together, the ChIP data and SURVEYOR results show that a site within closed chromatin (site sg025) can remain accessible to Cas9/sgRNA binding and DNA editing. The results also reveal that Cas9 editing can be blocked in the presence of Cas9/sgRNA-binding (site sg034). Downstream steps in INDEL formation, such as DNA cleavage or error-prone repair, may be reduced near compacted chromatin.

Next, we investigated whether changes in chromatin states (illustrated in Figure 6a) could enhance or restore Cas9-mediated editing. In order to induce a hyperactive expression state, we transfected Luc14 cells with a plasmid that expressed the strong transcriptional-activator Gal4-p65. We exposed the *luciferase* transgene to Cas9 editing by co-transfecting these cells with the Cas9/sg034 plasmid. Gal4-p65 induced luciferase expression approximately 6-fold compared to the basal expression level in Luc14 cells (Figure 6b). SURVEYOR showed reduced INDEL formation at hyperactivated *luciferase* compared to the basal level (p = 0.018), suggesting that dynamic chromatin remodeling through transcriptional activation or competition with transcription factors interferes with Cas9-mediated INDEL formation. Next, we investigated Cas9 efficiency after restoring the active state from a previously silenced state. GAL4EED +dox cells were treated with anti-SuZ12 siRNA to disrupt PRC2 accumulation, or Gal4-p65 to enhance *luciferase* expression. SuZ12, a member of the PRC2 complex, is required for maintaining the H3K27me3 mark^15^. Anti-SuZ12 stimulated partial reactivation compared to the basal expression state and showed full recovery of Cas9 editing efficiency compared to the fully silenced state (p = 0.045) (Figure 6b and c). Gal4-p65-treated GAL4EED +dox cells showed full reactivation of *luciferase* compared to the basal expression level and partially restored the frequency of INDEL formation (Figure 6b and c). These results suggest that artificial restoration of gene expression is an effective approach for enhancing Cas9-mediated editing at a target gene.

**Figure 6.**
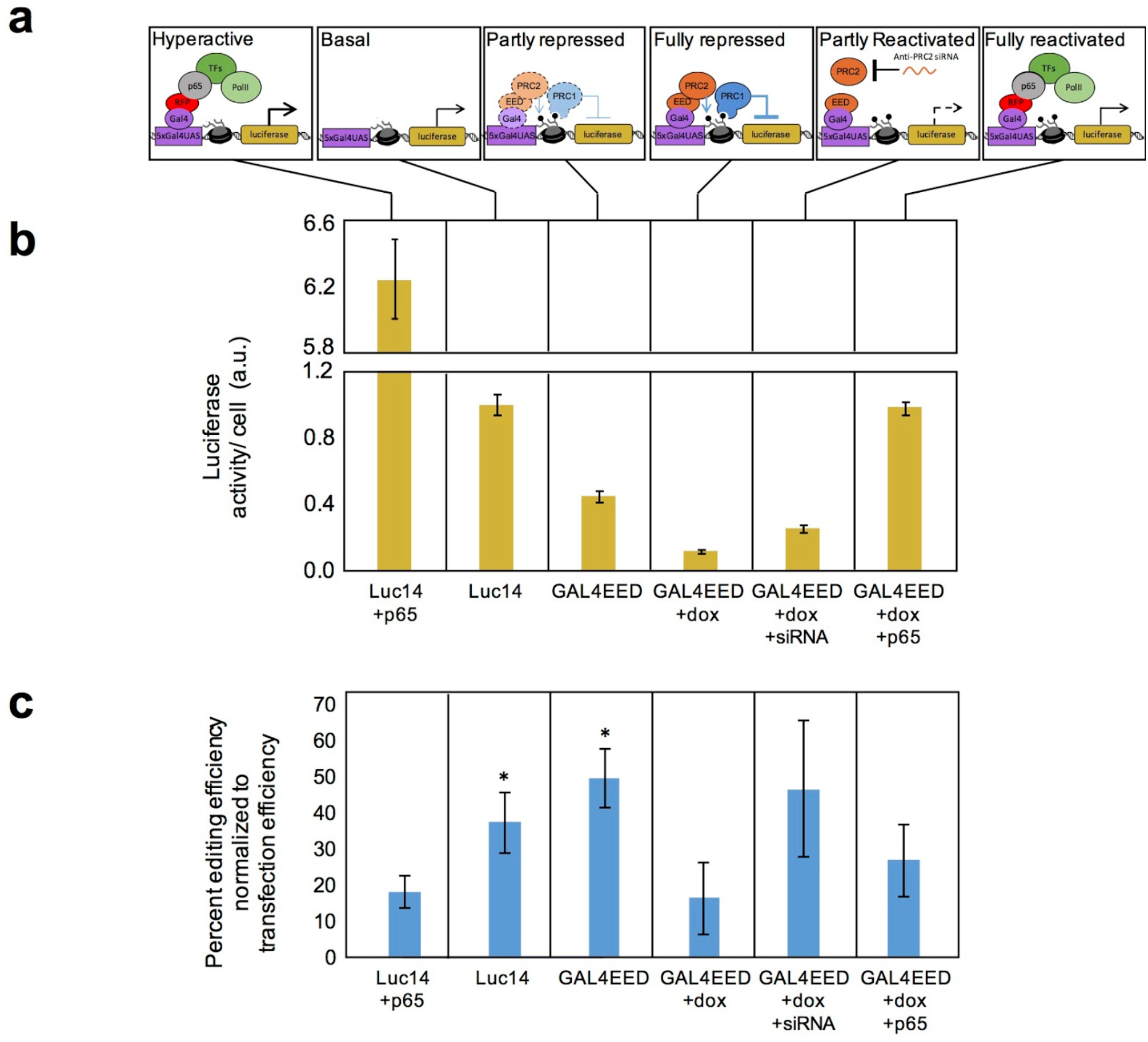
Dynamic regulatory states impact Cas9-mediated editing at the *luciferase* transgene. (a) Illustration of the *luciferase* transgene in the basal expression state and in different, artificially-regulated states. (b) Background-subtracted Luciferase expression levels per cell were measured 96 hours after dox treatment (GAL4EED +dox), immediately before transfection with Gal4-p65 plasmid DNA (+p65), or mock-transfection (vehicle only). Luciferase expression was measured in siRNA-treated cells 336 hours after transfection. a.u.: arbitrary units. (c) Editing efficiency for Cas9/sg034 was determined by SURVEYOR assays. Editing was reduced in the hyperactive expression state (Luc14 +p65) compared to the Luc14 basal state (*p = 0.018) and the GAL4EED partially repressed state (*p = 0.004). Reversal of the closed state via siRNA treatment (GAL4EED +dox +siRNA) was accompanied by an increase in luciferase expression and editing efficiency (p = 0.045) compared to the fully repressed state. Editing efficiencies for Luc14, GAL4EED, and GAL4EED +dox shown here are the same data shown in Figure 3c.

In conclusion, our results provide direct evidence that at a single genomic locus, Polycomb-mediated chromatin structure impairs Cas9-mediated DNA editing at specific sites. Specifically, in the fully repressed chromatin state, the sg034 target site is accessible to Cas9/sgRNA binding, but INDEL formation is reduced. Non-template-guided Cas9-mediated editing requires three steps: Cas9/sgRNA-target binding, endonuclease activity, and INDEL formation through error-prone NHEJ-mediated repair. Others have investigated the Cas9/sgRNA-target binding step using dCas9 and found off-target binding to be reduced at sites of closed chromatin^8,12,13^ While these data suggest possible interference at this step, the impact of closed chromatin on Cas9/sgRNA-DNA interactions at an on-target, specific binding was not determined. Recent work suggests that the nucleosome core particle directly contributes to Cas9 interference. Nuclease activity on naked DNA confirmed that Cas9/sgRNA is active *in vitro*^24^. Studies with reconstituted chromatin templates demonstrated that nucleosome-occupied regions are blocked from Cas9 binding^25,26^. We used inducible facultative chromatin in live HEK293 GAL4EED cells and ChIP-qPCR to compare dCas9 occupancy at *luciferase* in the open vs. closed chromatin state and observed that Cas9 protein binding is not reduced in the closed state. Therefore, at the transgenic locus analyzed here, chromatin may specifically interfere with Cas9 endonuclease activity, error-prone repair, or both. This conclusion is consistent with previous evidence that suggests that NHEJ may be slowed or inhibited in closed chromatin^27,28^. Taken together, these findings support a model for Cas9 inhibition where chromatin compaction and nucleosome occupancy disrupt Cas9-mediated editing at steps that follow Cas9/sgRNA binding.

Our findings identify repressive chromatin as a critical barrier to efficient Cas9-mediated editing in mammalian cells. In high-throughput applications, such as generating knockout libraries for model organisms, many target sites may be located in closed chromatin in certain cell types. Many sgRNAs may be prone to high failure rates, thus trial-and-error to achieve gene editing may be impractical. Methods for opening closed chromatin, such as transcription activators^29^, and inhibitors of heterochromatin proteins^30,31^, might enhance Cas9 editing efficiency. We observed that treating cells with siRNA against the silencing protein SuZ12 led to full recovery of editing efficiency, comparable to the basal expression state (Figure 6). Therefore, an effective general strategy for restoring an open, Cas9-accessible state at chromatin-regulated target sites may be siRNA-mediated chromatin inhibition. Our findings suggest that an effective strategy to increase Cas9 activity at closed chromatin should effectively induce open chromatin without causing interference by blocking Cas9’s access to the DNA target site. Our study with a switchable Polycomb chromatin system^15^ opens new and impactful avenues to define the role of eukaryotic chromatin in Cas9-mediated genome editing and to discover new strategies for enhancing Cas9 function in the context of the human genome.

## Methods

### Plasmid DNA construction

In order to determine transfection efficiencies using flow cytometry, we modified the vectors pX330-U6-Chimeric_BB-CBh-hSpCas9^4^ (a gift from Feng Zhang, Addgene plasmid #42230) and pX330A_dCas9-1×4 [Nakagawa 2015] (a gift from Takashi Yamamoto, Addgene plasmid #63598) using the T2A peptide skipping sequence to express EGFP from the same mRNA transcript as the Cas9 protein. PX330 or pX330A and the gBlock Gene Fragment (Integrative DNA Technologies) FseINLS-T2A-EGFP-EcoRI containing EGFP were cut with FseI (New England BioLabs) and FastDigest EcoRI (ThermoFisher Scientific) and ligated using T4 DNA Ligase (New England BioLabs). We named this new vector pU6-(BbsI)_CBh-Cas9-T2A-EGFP (DNASU UnSC00746685). SgRNAs used in the study (Supplemental Table 1) were designed using the CRISPR design tool at crispr.mit.edu9. DNA oligos were synthesized with the overhangs for cloning into pX330g or pX330g_dCas9 (Integrative DNA Technology). Drop-in of sgRNAs followed the cloning protocol described in Cong *et al.* 2013^4^. The Gal4-p65 fusion was expressed from plasmid CMV-Gal4p65_MV1 (DNASU UnSC00746686). Annotated sequences of the plasmids used in this study are available online (https://benchling.com/hayneslab/f_/V1mVw1Lp-chromatin-crispr-interference/).

### Cell culturing and transfections

The Luc14 cell line carries a *firefly luciferase* gene (*Gal4UAS-TK-luciferase*), which is stably integrated into the genome of HEK293 cells^14^. A second cell line, GAL4EED, contains the *firefly luciferase* gene (*Gal4UAS-TK-luciferase*) as well as a *TetO-CMV-Gal4EED* transgene, which carries a Gal4 DNAbinding motif (Gal4) fused to an embryonic ectoderm development (EED) open reading frame driven by TetO-CMV promoter^14^. Expression of the *Gal4EED* fusion protein-encoding sequence is silenced by a Tetracycline repressor (TetR) (Figure 1a). The cell line also contains a puromycin (puro)-inducible anti-Gal4EED miRNA to counter leaky transcription of *TetO-CMV-Gal4EED.* The removal of puro and addition of doxycycline (dox) to cultured GAL4EED cells releases the TetR protein from *TetO-CMV-Gal4EED* and allows the expression of Gal4EED. Gal4EED binds to the Gal4UAS site and switches the chromatin state at *luciferase* from active to silenced through accumulation of PRC (Figure 1B, Hansen et al., 2008^15^, and Haynes et al., 2011^28^).

Cells were grown in Gibco DMEM high glucose 1X (Life Technologies) with 10% Tet-free Fetal Bovine Serum (FBS) (Omega Scientific), 0.5 µg/mL puro and 1% penicillin streptomycin (ATCC) at 37°C in a humidified 5% CO_2_ incubator. At time 0 hours, puromycin was removed and GAL4EED cells were induced with doxycyclin (dox) (Santa Cruz Biotechnology) at 1 µg/mL. At 72 hours, Luc14 cells and dox-induced GAL4EED cells were seeded in 12-well plates such that at 96 hours, cells reached 90% confluency. One well from each cell type was collected for Luciferase assay (see below). The remaining wells were used for lipid-mediated transfection. For pX330g/sgRNA transfections, 0.5 µg plasmid was used / well, 3 µL Lipofectamine LTX, and 1 µL Plus Reagent (Life Technologies) per manufacturer’s protocol. For pX330g/sgRNA transfections, 0.5 µg plasmid was used / well, 3 µL Lipofectamine LTX, and 1 µL Plus Reagent (Life Technologies) per manufacturer’s protocol. At 144 hours, cells were collected to determine transfection efficiency by flow cytometry and editing efficiency by SURVEYOR Assay (see below).

For p65 CMV-Gal4p65_MV1 transfections, 0.5 µg of each plasmid was used / well, 3 µL Lipofectamine LTX, and 1 µL Plus Reagent (Life Technologies) per manufacturer’s protocol. At 144 hours, cells were collected for downstream analyses, including determining transfection efficiency, and performing Luciferase Assay and SURVEYOR Assay (described below).

### Luciferase assay

Cells were washed with 0.5 mL PBS (Irvine Scientific), detached with 0.2 mL trypsin (Life Technologies), collected with the addition of 0.5 mL DMEM, and spun at 100 RCF. Supernatant was aspirated and the cell pellet was resuspended in 2 mL of FACS buffer (PBS with 1% FBS) and filtered using 35 µm cell strainers (Electron Microscopy Sciences). 20 µL of cells + FACS buffer were counted and gated using the BD Accuri C6 Flow Cytometer and software (BD Biosciences). The luciferase assay was performed using Steady-Luc Firefly HTS Assay Kit (Biotium) according to manufacturer’s protocol. Briefly, 100 µL of cells in FACS buffer from each sample were added to three wells of a Corning and Costar 96-well Cell Culture Plate, black, clear bottom (Bioexpress). One hundred μL of Luciferase working solution was added to each well. Three wells with FACS buffer (without cells) plus Luciferase working solution were read and the highest measured value of the three wells was used as background signal. The plate was incubated for 5 minutes with orbital shaking and luminescence was read using a Synergy H1 Multi-Mode Reader (Biotek). Luciferase expression per cell was calculated as:

> Sample Luciferase per cell = [Sample Luciferase signal] - background signal /[cell count * (100 µL / 20 µL)]

### SURVEYOR assay

Genomic DNA was extracted using a QIAamp DNA Mini Kit (Qiagen). Cas9/sgRNA target DNA was amplified using Phusion polymerase first with external primers P197 5’-gctcactcattaggcacccc and P198 5’-ggcgttggtcgcttccggat. PCR products were diluted 1:1,000 in water. Nested PCR was performed using primers flanking the Cas9/sgRNA target site (Supplemental Tables 2 and 3). An annotated map of primers is available online (UAS-TK-luc HEK293, https://benchling.com/hayneslab/f_/V1mVw1Lp-chromatin-crispr-interference/). PCR products were purified (GenElute PCR Clean-Up, Sigma) and SURVEYOR assay (IDT) was performed according to the manufacturer’s protocol. Briefly, 400 ng of PCR product was mixed with 1.5 µL of annealing buffer (10 mM Tris-HCl (pH 8.8), 1.5 mM MgCl2, and 50 mM KCl), and brought to 15 µL with water. PCR products were melted and reannealed and digested with 1 µL SURVEYOR enzyme and 1 µL Enhancer for 1 hour at 42°C. The concentrations of fragments in each sample were measured on an Agilent 2100 Bioanalyzer. The ratio of uncut wildtype (WT) to cut heteroduplex DNA fragments (HDlarge, HDsmall) was calculated as:

> Percent editing efficiency = 100 * [HDlarge + HDsmall] / [ HDlarge + HDsmall + WT]

Genomic DNA from untreated Luc14 and GAL4EED cells and genomic DNA from Cas9/sgRNA treated cells lacking S-nuclease treatment were used as controls to distinguish background noise from actual cut heteroduplex DNA fragments.

The editing efficiencies shown in Figures 3 and 6 were normalized by transfection efficiency. At 72 hours post-transfection (12-well plates, ∼0.4×10^6^ cells/well), cells were washed with 0.5 mL PBS (Irvine Scientific), detached with 0.2 mL trypsin (Life Technologies), collected with 0.5 mL DMEM, and spun at 100 x g for 5 minutes. Supernatant was aspirated and the cell pellet was resuspended in 2 mL of FACS buffer (1% FBS in 1xPBS) and filtered using 35 µm cell strainers (Electron Microscopy Sciences). Live cells were gated based on forward and side scatter using the BD Accuri C6 Flow Cytometer and software (BD Biosciences). Data was analyzed using FlowJo software. Fluorescence intensity for ten thousand live cells, detected with settings for GFP (488 nm laser, 533/30 filters), was plotted against cell count. The GFP-expression threshold was determined using non-GFP expressing HEK293 cells. Transfection efficiency was calculated as the percent of cells GFP-positive cells in the total live cell population. This value was used to normalize editing efficiencies to cells containing the Cas9/sgRNA plasmid:

> Percent editing efficiency normalized to transfection efficiency = [Percent editing efficiency] / [transfection efficiency]

### Statistical analyses

For analyses of SURVEYOR Assay data, standard deviations were calculated for n=3 biological replicates. The difference of means for GAL4EED and GAL4EED +dox was calculated using the two sample, one-tailed Student’s t-test with a confidence of 97.5% for 2 degrees of freedom and a test statistic of t_(.025,2)_ = 4.30265.

### Mutant clone library

The sg034 target region was PCR amplified from genomic DNA and prepared as described above (see “SURVEYOR assay”). Thirty ng (0.072 pmols) of each blunt ∼630 bp PCR product was ligated with linear pJET1.2 vector (0.05 pmol ends) in a 10 µL reaction following the manufacturer’s protocol (CloneJET PCR Cloning Kit, Life Technologies) with the following modifications: 1x ligation buffer (Roche), T4 DNA ligase (New England Biolabs). Reactions were incubated at room temperature (25°C) for 5 min, mixed with 50 µL of thawed Turbo competent DH5-alpha *E. coli* (New England Biolabs), and incubated on ice for 5 minutes. Transformed cells were plated directly on pre-warmed LB agar (100 µg/mL ampicillin) and incubated overnight at 37°C to grow colonies. Liquid cultures (200 µL LB broth, 100 µg/mL ampicillin) were inoculated in deep round-bottom 96-well plates with single colonies collected via sterile, disposable pipette tips. Plasmid DNA was purified using the Montage Plasmid Miniprep HTS 96 Kit (Millipore). Sanger sequencing was performed using primer P163 (5’-caaaccccgcccagcgtctt). The sequence data were aligned to the pUAS-TK-luc reference sequence in Benchling using MAFFT^32^. Sequence variants were binned manually and counted using Excel software.

### Cross-linked chromatin immunoprecipitation (ChIP)

Luc14 or GAL4EED +dox cells were electroporated with plasmid pX330g_dCas9/sg025 or pX330g_dCas9/sg034 using the Neon Transfection System (Invitrogen) following manufacturers protocols. Electroporation settings were as follows: 100 µL tip, Pulse voltage 1,100 V, pulse width 20 ms, 2 pulses, with cell density 5 × 10∧7 cells/mL. Two transfections were plated into each of three 15 cm plates for each cell type for each of three replicates. Transfected cells were grown at 37°C for 72 hours, collected by trypsin-treatment, and incubated with 20 ml of 1% formaldehyde (Thermo Fisher Scientific) /1× Dulbecco’s PBS for 10 min at room temperature. Cross-linking was quenched with 125 mM glycine for 5 min. Cross-linked cells were washed twice with cold 1X PBS buffer + Pierce Protease Inhibitors (Thermo Fisher Scientific) for 5 minutes with shaking. Cells were washed again with 1X PBS buffer + Pierce Protease Inhibitors without shaking. Cells were spun at 1000 rpm for 5 min between each wash step.

Cells were diluted in ChIP Lysis Buffer [1% sodium dodecyl sulfate (SDS) (Sigma), 10 mM ethylenediaminetetracacetic acid (EDTA) (Fisher Scientific), 50 mM Tris-HCl pH 8.1 (Sigma)] to a concentration of approximately 4×10^7^ cell/mL. Samples were split into approximately 100 µL per tube and disrupted using a Qsonica Q700A Sonicator with a 5.5″ Cup Horn. Sonicated chromatin was spun at ∼10,000 × g for 10 minutes at 4°C to remove impurities and then flash frozen at −80°C.

DNA from 10 µL of unsonicated control and 10 µL sonicated chromatin was incubated at 65°C overnight (<16 hrs) in 0.1 M NaCl, then 30 minutes at 37°C with 10 ug of RNase A (Sigma), and then for 2 hours at 62°C with 10 ug of Proteinase K (Qiagen). DNA was resolved via electrophoresis to confirm ∼500-bp fragments.

For immunoprecipitations, 50 ug of each chromatin preparation was diluted to 1 mL in dilution buffer [1% Triton X-100 (Santa Cruz Biotech), 2 mM EDTA, 150 mM sodium chloride (NaCl) (Sigma), 20 mM Tris-HCl, pH. 8.0]. Chromatin was pre-cleared for 3 hours at 4°C with nutation with 20 ul of washed [3x PBS buffer + BSA (5 mg/mL) (Sigma)] Magna ChIP Protein A + G (Millipore). Fifty ul (20%) of pre-cleared chromatin was frozen for input control. Chromatin from non-transfected Luc14 and GAL4EED cells was incubated with 5 µg of rabbit anti-H3K27me3 07-449 (Millipore) or 5 µg of rabbit IgG (Cell Signaling 27295) at 4°C for 12 hours with nutation. Chromatin from dCas/CRISPR plasmid-transfected cells was incubated with 5 µg mouse anti-FLAG M2 Antibody (F1804, Sigma) or 5 µg of rabbit IgG (Cell Signaling 27295) at 4°C for 12 hours with nutation. Antibody-chromatin samples were incubated with 20 µL Magna ChIP Protein A + G beads for 3 h at 4 °C with nutation. Chromatin-antibody-bead complexes were washed 6 times with 10 minute rotating incubations with RIPA buffer [50 mM HEPES pH 7.6 (Thermo Fisher Scientific), 1 mM EDTA, 0.7% Sodium-Deoxycholate (Sigma), 1% IGEPAL CA-630 (Sigma), 0.5 M LICl (Sigma)] and two times with 5 minutes rotating incubations with tris-EDTA pH 7.6 (Sigma). Washed chromatinantibody-bead complexes were resuspended in 100 ul of elution buffer [1% SDS, 0.1 M sodium bicarbonate (Sigma), 0.1 M NaCl]. Fifty μL inputs were brought up to 100 µL in elution buffer. IPs and inputs were nutated for 30 minutes at room temperature then incubated at 65°C for 9 hours to reverse cross-linking. Samples were treated for 30 minutes at 37°C with 10 ug of RNase A and then for 2 hours at 62°C with 10 ug of Proteinase K. DNA from IPs and inputs was purified with Genelute PCR Cleanup Kit (Sigma) and eluted in 50 µL nuclease-free water.

### Real-time Quantitative PCR of ChIP-enriched DNA

Real-time quantitative PCR reactions (15 µl each) contained SYBR Green master mix, 2 µl of immunoprecipitated (IP), IgG-IP, or input template DNA, and 2.25 pmol of primers. Input Cp values were adjusted by subtracting log2(20) from each input Cp, as 50 µL was taken from 1 mL total chromatin or 1/20. % IP DNA bound was calculated as 100 × 2^[Ct input − Ct IP]^. % IgG-IP bound (100 × 2^[Ct input − Ct IgG-IP]^ was subtracted from % IP DNA bound to calculate “% IP DNA enrichment relative to IgG-IP.”

Primer sequences for site 1 were

5’-cgaccctgcataagcttgcc (forward)
5’-ccgcgtacgtgatgttcacc (reverse).
Primer sequences for site 2 were

5’-gcgcccgcgaacgacattta (forward)
5’-gagatgtgacgaacgtgtac (reverse).
Primer sequences for site 3 were

5’-gcgcccgcgaacgacattta (forward)
5’-gagatgtgacgaacgtgtac (reverse).
Primer sequences for site 4 were

5’-ttgtaccagagtcctttgatcg (forward)
5’-ccgtgatggaatggaacaac (reverse).
Primer sequences for GAPDH were

5’-tactagcggttttacgggcg (forward)
5’-tcgaacaggaggagcagagagcga (reverse).

### siRNA transfections

At time 0 hours, puromycin (puro) was removed and GAL4EED cells were induced with doxycycline (dox, Santa Cruz Biotechnology) at 1 µg/mL. At 96 hours, dox was removed and media with puro was used. Cells were seeded in 12-well plates such that at 96 hours, cells were at 50% confluency. One well was collected for Luciferase assay (see above). The remaining wells were used for lipid-mediated transfection with 2.5 µL of 20 µM anti-SUZ12 siRNA duplex (Dharmacon) / well and 1.5 µL Oligofectamine (Life Technologies) per manufacturer’s protocol. siRNA sequence used^15^:

Sense: 5’ A.A.G.C.U.G.U.U.A.C.C.A.A.G.C.U.C.C.G.U.G.U.U 3′
Antisense: 5′ C.A.C.G.G.A.G.C.U.U.G.G.U.A.A.C.A.G.C.U.U.U.U 3′

Mock transfected cells (water used in place of siRNA duplex) were used as a control. At 144 hours, siRNA and mock transfections were repeated. At 264 hours, siRNA and mock transfected cells were transfected with Cas9/sg034 (three experimental replicates) (see above). At 336 hours cells were collected to determine luciferase expression, transfection efficiency, and editing efficiency (see above).

## Supporting Information Available

Supplementary Figure S1, Chromatin mapping data show enrichment of H3K27me3 across the Cas9-targeted region at *luciferase*; Supplementary Table 1, List of sgRNAs; Supplementary Table 2, List of primers for nested PCR of genomic DNA from Cas9/sgRNA treated samples; Supplementary Table 3, List of primers with their sequences. This material is available free of charge via the Internet at http://pubs.acs.org. Annotated sequences of the plasmids used in this study are available online (https://benchling.com/hayneslab/f_/V1mVw1Lp-chromatin-crisprinterference/). Plasmids have been deposited in the DNASU repository and are available under accession codes UnSC00746685 (http://dnasu.org/DNASU/GetCloneDetail.do?cloneid=746685) and UnSC00746686 (http://dnasu.org/DNASU/GetCloneDetail.do?cloneid=746686).

## Abbreviations

CRISPR: Clustered regularly-interspaced palindromic repeats
PRC: Polycomb Repressive Complex
sgRNA: short guide RNA
INDEL: insertion and deletion

## Author Information

Correspondence and requests for materials should be addressed to Rene Daer, Arizona State University 501 E. Tyler Mall, ECG 344A, Tempe AZ 85287, United States. The authors declare they have no competing financial interests.

## Acknowledgement

The authors thank J. Steel (DNASU) for DNA sequencing services and the 2014 Cold Spring Harbor Synthetic Biology Summer Course for supporting our development of this study. The authors thank F. Ceroni, J. Eisenberg, C. Barrett, D. Nyer, D. Vargas, and K. Timms for critical review of the manuscript. KAH is supported by the NIH NCI (K01CA188164) and the NSF/Synberc (EEC 0540879). RMD is supported by the ARCS Foundation and NSF CBET (1404084).

## Author Contributions

RMD performed DNA construction, cell culturing and assays, and SURVEYOR assay. The clone library was generated by KAH and sequenced by DNASU. RMD and JPC performed ChIP assays. All authors analyzed the data and composed the figures and text.

